# Inferring drift, genetic differentiation, and admixture graphs from low-depth sequencing data

**DOI:** 10.1101/2024.01.29.577762

**Authors:** Malthe Sebro Rasmussen, Carsten Wiuf, Anders Albrechtsen

## Abstract

A number of popular methods for inferring the evolutionary relationship between populations require essentially two components: First, they require estimates of *f*_2_-statistics, or some quantity that is a linear combination of these. Second, they require estimates of the variability of the statistic in question. Examples of methods in this class include qpGraph and TreeMix.

It is known, however, that these statistics are biased when based on genotype calls at low depth. Moreover, as we show, this leads to downstream inference of significantly distorted trees. To solve this problem, we demonstrate how to accurately and efficiently compute a broad class of statistics from low-depth whole-genome sequencing data, including estimates of their standard errors, by using the site frequency spectrum. In particular, we focus on *f*_2_ and the sample covariance of allele frequencies to show how this method leads to accurate estimate of drift when fitting trees using qpGraph and TreeMix with low-depth data. However, the same considerations lead to uncertainty estimates for a variety of other statistics, including heterozygosity, kinship estimates (e.g. King), and quantities relating to genetic differentiation such as *F*_st_ and *D*_*xy*_.

## Introduction

Inference and analysis in population genetics using sequencing data typically starts with variant and genotype calling. However, it has become clear that when sequencing at lower depth, where the uncertainty of the call is greater, this significantly biases a wide range of downstream analyses [1–3]. At the same time, low-depth data may be the only available option, as may be the case when working with ancient DNA [4, 5]. More broadly, sequencing at lower depth is beneficial to achieve larger sample sizes at a fixed sequencing cost. In fact, assuming that the use of low depth sequencing can be measured as citations per year of the low depth sequencing software ANGSD [6], then usage has been monotonically increasing each year. Therefore, the ability to perform sound inference based on low-depth data remains important despite decreasing costs of sequencing.

To properly account for genotype uncertainty, a very large number of methods have been tailor-made to low-depth data [3]. Creating, testing, and implementing such methods is time-consuming for developers, and also require practitioners to keep up with a bifurcated ecosystem with a different set of tools for high and low-depth data. However, a broad class of statistics (and associated methods) can be re-written in terms of the site frequency spectrum (SFS). This turns out to be helpful, since whereas individual genotypes cannot be reliably called at low depth, we have good methods available for the SFS in this context [1, 7–9]. In practice, that implies that any statistic or method that can be reformulated in terms of the SFS is immediately amenable to use with low-depth data. A number of existing methods are instances of this general strategy [10, 11], which we formalise below.

However, a barrier remains when not only some SFS-derived statistic is required, but also an estimate of its variability. Such an estimate is often useful in its own right to aid interpretation, but it may also be a requirement for certain procedures. Throughout, we focus on the examples of qpGraph (in the ADMIXTOOLS [12, 13] suite) and TreeMix [14], which fit trees based on *f*-statistics and sample allele frequency covariance, respectively. In both cases, an estimate of the variability is required. The end effect is that low-depth sequencing studies must currently run such tools with genotype calls, despite the knowledge that this likely introduces bias, or refrain from their usage altogether.

Our contribution in this article, then, can be seen as three-hold: First, we outline how *f*-statistics and sample covariance can be expressed as functions of the two-dimensional SFS to be used in a low-depth setting. Second, we provide a pragmatic approach to estimation of standard errors on SFS-derived statistics at the genome scale using a genomic jackknife. Finally, we show how this leads to significant improvements when fitting trees with qpGraph and TreeMix from low-depth data. We emphasise, though, that the underlying utility of having SFS-derived statistics with standard errors is more widely useful, and we provide other examples of this throughout.

## Methods

To begin, we outline the relevant background on the site frequency spectrum and derived statistics. At first, we will assume that genotypes can be accurately called, and we return below to the setting where this assumption cannot be made.

### Site frequency spectrum

Consider an index set of populations *𝒦* = {1, …, *K*}. We assume *N*_*k*_ diploid individuals have been sampled in the *k*th population, and sequencing data for *M* sites is shared across all samples.

Let *G*_*mnk*_ ∈ {0, 1, 2} be the genotype of individual *n* in population *k* at site *m* coded as the number of derived alleles observed at this position. Then we define the allele count

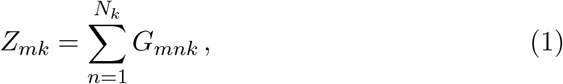

as the total number of derived alleles observed in sample *k* at site *m*. Note that *Z*_*mk*_ ∈ {0, …, 2*N*_*k*_} at any *m* so that, taken jointly, we have

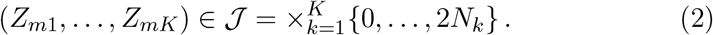

In other words, *𝒥* is the set of possible allele count configurations across populations *𝒦* at any site *m*.

Using this notation, we define the SFS for *𝒦* as the function *S* : *𝒥 →* [0, 1]:

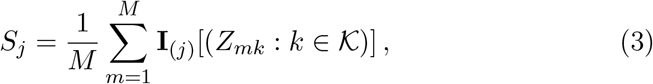

where **I**_*x*_[*P*] is the indicator function for a predicate *P* on *x*, such that *S*_*j*_ is the fraction of sites where the allele count configuration *j* ∈ *𝒥* is observed.

For example, suppose *𝒦* consists of just two populations. Then *S* can be thought of as a (2*N*_1_ + 1) *×* (2*N*_2_ + 1) matrix where *S*_{1,2}_ is the fraction of sites that are singletons in the first population and doubletons in the other.

For simplicity, we have taken the population sample set *𝒦* on *S* to be implicit, but often we are interested in the spectra for various subsets of the sample populations. Where necessary, we use the notation *S*(*𝒦* ^*′*^), *𝒦* ^*′*^ *⊆ 𝒦* to denote the SFS for a particular set of populations, and likewise *𝒥* (*𝒦* ^*′*^) for the corresponding allele count possibilities. To be clear, note also that we consider the SFS *S* to be a statistic of a sample, rather than as a population parameter.

### Derived statistics

The SFS in effect compresses all information about the joint sample allele frequencies in a set of samples. As a result, many statistics that are functions of sample allele frequencies (and sample sizes) can be expressed as functions of the SFS.

Specifically, let

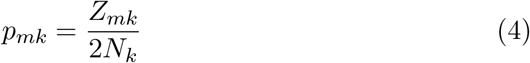

be the sample allele frequency for population *k* at site *m*, and let *T* be a statistic that is a site-wise average over a function *f* of joint sample allele frequencies (and sample sizes) in *𝒦*. Then, *T* is also a function *g* of *S* according to,

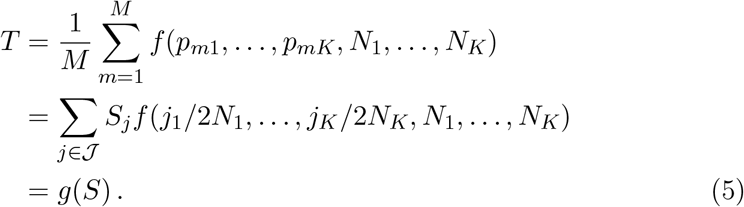

We refer to a such statistic *T* as a derived statistic of *S*. In the following, we illustrate this with the *f*-statistics and sample allele frequency covariance.

### *f*-statistics

The quantities *f*_2_, *f*_3_, and *f*_4_ [12, 15] relating two, three, and four populations, respectively, are informative of their evolutionary relationships in a variety of ways [16, 17]. Among other things, these so-called *f*-statistics are used for demographic inference in the ADMIXTOOLS framework [12, 13].

We understand these *f*-statistics not as statistics in the strict sense, but rather as unobserved quantities to be estimated. Specifically, let *k*_1_, *k*_2_, *k*_3_, *k*_4_ ∈ *𝒦*, and the standard estimators are given by,

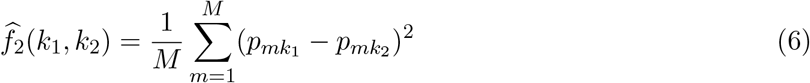

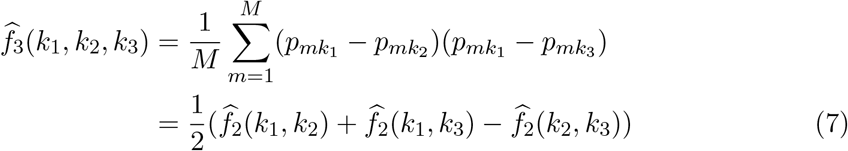

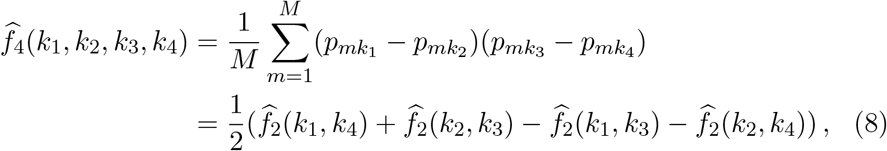

and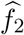 can be written as a function of the corresponding two-dimensional SFS,

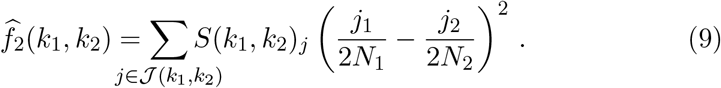

Likewise, 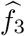 and 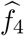 can be written as functions of the three and four-dimensional spectra, respectively. However, note that they can also be written as linear combinations of functions of multiple two-dimensional spectra. We return to this point below.

### Sample covariance

The covariance of allele frequencies is another parameter that is useful for population genetic inference. For instance, it serves as the basis of inference for TreeMix [14]. However, the expectation of allele frequencies is not generally known, so they use instead the mean population allele frequency. That is, for populations *k*_1_, *k*_2_ *∈ 𝒦*, define the sample covariance *k*_1_ and *k*_2_ with estimator

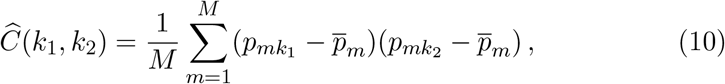

where

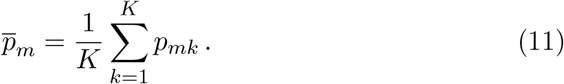

It is possible to write *Ĉ* as a function of the full, *K*-dimensional SFS. However, for reasons described below, we will make use of the fact that *Ĉ* can be written as follows [18]:

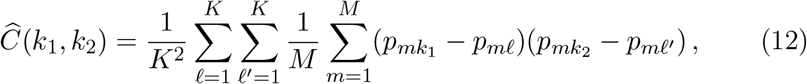

where every inner term is some 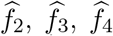 or trivially zero. Any of those terms can be expressed in terms of 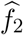 and it therefore follows that *Ĉ* can be expressed as a function of the full set of two-dimensional spectra for all combinations in *𝒦*.

### Variability

It may be desirable to calculate not only some statistic *T* from the SFS, but also some estimate of the variability of *T*. This may be useful in its own right for interpretation of results, and it may be required for further inference. For instance, both ADMIXTOOLS and TreeMix require an estimate of the standard error of the input 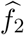 and *Ĉ* statistics, respectively, in order to run.

Typically, the relevant sense of variability is both technical as well as biological variability. That is, we are interested in how the statistic varies over the genome. For this purpose, a popular approach is to use a genomic jackknife to estimate standard error.

For the genomic jackknife, the data is split site-wise into *B* blocks *ℬ* = {*ℬ*_1_, …, *ℬ*_*B*_} so that blocks are non-overlapping, exhaustive, and all sites in *B*_*i*_ precede (in a genomic sense) *B*_*j*_ where *i < j*. Let *T* ^[*−i*]^ refer to the statistic *T* calculated from the data *ℬ \ ℬ*_*i*_, and let *T* be the statistic calculated on the full data set. Following [19], we then define

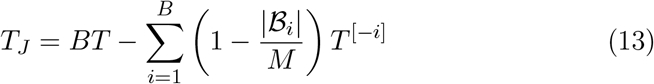

and

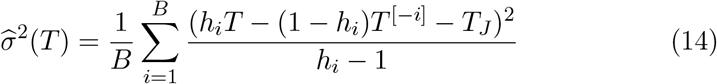

with *h*_*i*_ = *M/*|*ℬ*_*i*_|, is an estimate of the variance of *T*_*J*_, and is used in practice as an estimate of the variance of *T*.

We can estimate the variance of the SFS *S* with this procedure, as well as any statistic *T* derived from the SFS, since *T* ^[*−i*]^ = *g*(*S*^[*−i*]^) in the notation of eq. (5).

### Low-depth data

In summary, we have so far outlined a general approach to estimating a class of SFS-derived statistics together with their standard errors. In various forms, this approach is in wide use in the field today. The issue is that it does not work when applied to low-depth sequencing data. As a result, a whole class of methods are currently unavailable to researchers working with such data, or available only with significant caveats.

To back up, we have so far assumed that *S* is a statistic that can be calculated from a sample by straight-forward application of eq. (3) by simply tallying genotypes. With high-depth data, where genotypes can be called with high confidence, this is true for all practical purposes. However, with low-depth data, we must treat *S* as an unobserved quantity to be estimated. For this setting, we introduce the notation 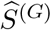 for the estimator of *S* based on eq. (3) using genotype calls. We can then be more specific with regards to the issue introduced above and note that 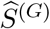 is known to be significantly biased [2, 7]. Further, this bias cannot be mitigated by filtering sites and is known to propagate to downstream statistics and inference [7, 9].

Instead, the preferred approach is to estimate *S* directly from genotype likelihoods without calling genotypes. Briefly, the genotype likelihood *p* (*X*_*mnk*_ | *G*_*mnk*_) corresponds to the probability of the sequencing data at site *m* for individual *n* in *k* given the genotype *G*_*mnk*_. The exact nature of sequencing data *X*_*mnk*_ is not important here, but may be thought of as read pile-ups, for instance. Instead, it suffices to note that genotype likelihoods are calculated from sequencing data as standard by a range of common bioinformatic tools [6, 20–22].

The typical method is to find the maximum-likelihood estimate of the site frequency spectrum:

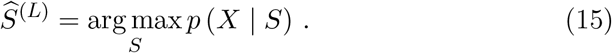

Eq. (15) often has no closed form solution, so an iterative optimisation scheme is employed, typically an EM algorithm. This approach has been shown to provide an approximately unbiased estimate of the SFS at low depths [2, 7, 9].

However, the situation is less straight-forward when trying to estimate the variability of 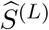 and derived statistics, With genotypes, we can carry out computations based on

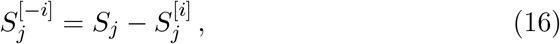

where 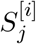 refers to the SFS calculated only on the block *B*_*i*_. Hence, we simply need all the 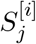 which can be found in linear time.

The same does not strictly hold for the an EM algorithm (or other iterative methods) for 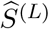 Of course, it is theoretically possible to simply directly estimate *S*^[*−i*]^ for all *i*, but the computational demands of this can be prohibitive at the genome scale: the runtime complexity of the standard EM algorithm for SFS estimation is 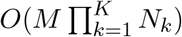 per iteration, so computation of all *S*^[*−i*]^s approaches 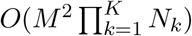 time per iteration as *B* increases, which is not feasible.

Instead, we make the assumption that eq. (16) will be approximately true for whole-genome sequencing data when *M* is large enough to pose a challenge and *B* takes on a typical low value (say, less than 100) so that each block contains in the tens or hundreds of megabases. We return to evaluate the implications of this in the results section below. Having made this assumption, only the global estimate and the block estimates are required, and these can be estimated in linear time for fixed populations.

In principle, this enables the same methods as those outlined in the previous sections to work with low-depth data. Nevertheless, since the run-time for SFS estimation is exponential in the number of populations involved, it remains infeasible to estimate spectra for more than two or three populations jointly at the genome scale. It is for this reason that we stated *Ĉ* as a combination of *f*-statistics in eq. (12): computing a full, *K*-dimensional SFS to calculate *Ĉ* will rarely be possible, but calculating *K*(*K* + 1)*/*2 two-dimensional spectra and the associated 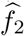 statistics typically will be.

### Implementation

We implemented the block SFS estimation as a sub-command of the winsfs software [9]. By default, the program splits the input into 50 blocks of approximately equal size, and estimates the spectrum for each of those in parallel using the standard EM algorithm. To speed up convergence, the estimation in each of the blocks is started using an estimate of the global SFS, which in turn is estimated using the main winsfs mode with default settings. Each block is run until the change in log-likelihood between successive iterations of the EM algorithm falls below 10^−8^ by default.

## Results

We evaluate the performance of this blocked SFS procedure for inference on low-depth data in the sections below.

### Simulations

To evaluate the validity of the proposed method and its assumptions, we tested the method on simulated sequencing data from a stepping stone demographic scenario simulated using `msprime` [23]. The simulated model consists of seven populations, which we refer to as A through G. Beginning with A, each population successively splits off from the previous, and immediately after the split undergoes a bottleneck before recovering its population size. In effect, this creates a cline of drift, where less drift has occurred on A and more drift has occurred on G. The basic model structure is shown in figure 1a.

**Figure 1:**
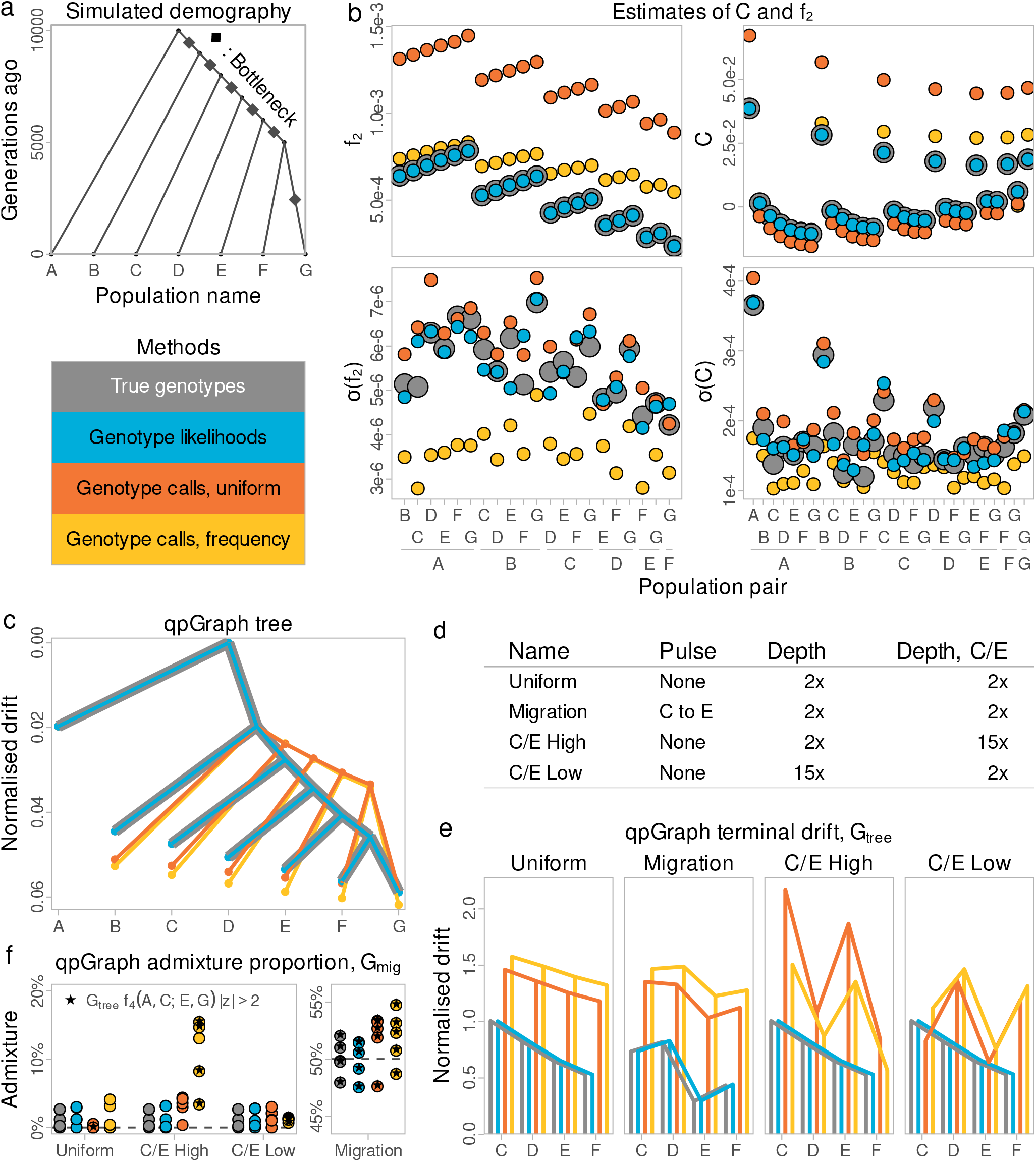
Simulation results. Lines and points for true genotypes are drawn larger to be visible underneath those for to genotype likelihoods. **(a)**: Overview of the simulated demographic history. **(b)**: 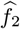 and *Ĉ* with standard errors. **(c)**: Drift estimated by qpGraph, normalised by the drift on the outgroup. **(d)**: Overview of the alternative models. **(e)**: Terminal drift estimated by qpGraph on *G*_tree_ for populations C, D, E, F for all simulations, normalised as in 1c. Drift between internal nodes not to scale. **(e)**: Admixture proportion estimated by qpGraph on *G*_mig_ for all simulations. Five independent replicates shown with true admixture proportion overlaid. Replicates are marked if qpGraph rejects (i.e. *z >* 2) the fit of *f*_4_(A, C; E, G) in *G*_tree_.

We simulated 22 independent 10 Mb chromosomes using mutation and uniform recombination rates of 2.5 · 10^−8^ and 10^−8^ per generation per site, respectively. For simplicity, we ignored multiallelic sites that arose during simulation. Based on the remaining data, we calculated all true one and two-dimensional spectra as well as spectra for 50 blocks of equal sizes using the `sfs` CLI utility.

Using the true genotypes, we imitated the effects of sequencing by using vcfgl to calculate genotype likelihoods based on a Poisson distributed depth with mean 2 x and an error rate of 0.2 %. Genotype likelihoods were simulated for all sites in the 220 Mb—as opposed only those that were polymorphic. From these, we used `angsd` [6] to compute site allele frequency likelihoods as input for `winsfs` to estimate the global and block spectra with default settings, thereby splitting the data into 50 equally sized blocks.

As a point of comparison, we also used the simulated genotype likelihoods to re-call genotypes using `angsd`. Here, the most common approach would be to use a prior on the genotypes, but using a uniform prior may often be the better choice when forced to call genotypes at low depth [24]. Therefore, we called genotypes either based on the maximum genotype likelihoods or with a prior based on the site-wise allele frequency estimated from the genotype likelihoods using an EM algorithm [6]. From the called genotypes, we calculated the SFS in the same 50 genomic intervals. For a given SFS or block SFS, we ignored sites with any missing genotype calls in the populations involved.

In review, this leaves us with four different methods for producing the SFS and associated blocks. For each of the four, we calculate and compare derived statistics and standard errors. To summarise, we introduce the following terminology:

1. True genotypes; we will refer to this as the truth, so that e.g. the true 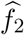 efer to 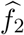 based on the SFS using true genotypes.
2. Genotype likelihoods, using EM algorithm implemented in `winsfs`; we refer to this as the GL method.
3. Genotype calls with a uniform prior.
4. Genotype calls with a frequency prior.

It is these four methods that are compared in figure 1 as described below.

### *f*-statistics and sample covariance

For each method and for all population pairs, we calculated 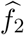 from the global SFS and all blocks. From the block estimates, we estimated the standard error 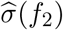 with a genomic jackknife as described. For the genotype call methods, where missingness cause blocks to have uneven sizes, these were weighted by the number of sites per block.

Using the *f*_2_ estimates, we calculated 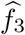 and 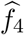 and used all *f*-statistic estimates to calculate *Ĉ*. Since 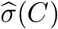 is computed as a linear combination of statistics with various amount of missingness, we used an unweighted jackknife in all cases. The resulting estimates are shown in figure 1b for all population pairs and for all four methods—note that the inclusion of monomorphic sites in the SFS cause the estimates to come out at a lower scale than when using only SNPs.

We observe that the GL method produces estimates 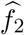 and *Ĉ* that are essentially identical to the true genotypes. In contrast, genotype calling leads to an upwards bias in 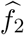_2_ that looks like additional drift between population pairs. For the same reason, *Ĉ* based on genotypes calls is inflated on the diagonal (i.e. for *Ĉ* (*k*_*i*_, *k*_*i*_)) and elsewhere underestimated.

For the standard errors, the GL method is not identical to the estimate from the true genotypes, but appear to be approximately unbiased relative to these. That makes sense assuming that the technical variability induced by the effects of sequencing is dominated almost entirely by biological variability when using the genotype likelihood based method.

To verify that these results are not contingent on the particulars of a single simulation, we repeated the entire workflow for five differently seeded replicates. The results (figures S1 and S2) confirm that the described patterns are consistent across replicates.

Based on the raw values of the estimates of *f*_2_ and *C* and their standard errors, the GL method performs well both relative to the true genotypes and in comparison to the genotype calls. However, the impact of biased estimates on the estimates of interest is not obvious. On the one hand, it is possible that the small differences between the GL estimates and the true genotypes matter; on the other hand, it may be the bias from genotype calling could be insignificant. In particular, looking at figure 1b, someone might suspect that genotype calling with a uniform prior simply leads to a rescaling of 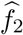 which would be inconsequential for downstream inference. To investigate, we proceed with further inference based on the raw estimates.

### Trees

Having estimates of *f*_2_ and *C* with standard errors enable methods like qpGraph in ADMIXTOOLS2 [13] and TreeMix [14]. As noted, *f*_2_ estimates with monomorphic sites are on a lower scale than when based solely on variable sites, which is the standard input for the above methods. To estimate drift on the same scale, we re-scaled all values of 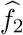 assuming 1 % polymorphic sites and re-calculated *Ĉ*s and standard errors before running these tools. This only changes the unit of the branch lengths and makes them comparable between methods without further normalization.

To begin, we used qpgraph to fit the drift on the simulated tree topology. The interface in ADMIXTOOLS2 does not allow user-calculated standard errors, so we used the raw 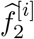 block estimates and let `admixtools` run an unweighted genomic jackknife. We are less interested in the absolute scale of drift than the relative drift within the tree, and therefore, to facilitate comparison between the methods, we also show the trees with an additional normalisation where the entire tree for each method is normalized by their drift on the outgroup (i.e. the branch length from the root to population A). The resulting tree are shown in figure 1c, and the unnormalised version in figure S4. We see that the GL method is again essentially equivalent to the true genotypes. Calling genotypes, regardless of the prior, however, generally leads to under-estimating drift on the internal branches and over-estimating the drift on the terminal branches. Taken together, this has the effect of significantly distorting the shape of the tree.

To run TreeMix, we modified the source code of `treemix` to allow running the core functionality directly from *Ĉ* and 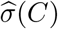A git patch file with the changes is available through the repository associated with this article. Using the modified program, we fit a tree with zero migrations and the true outgroup specified. The resulting drift estimates (figures S5 and S6) are essentially identical to those from qpGraph, emphasising the problem with genotype calling.

### Alternate simulations

So far, we have looked at simulations where the populations have the same mean depth of 2 x. However, issues with genotype calling are often exacerbated when different populations are sampled at varying depths. For instance, since errors in genotype calling can look like drift, calling genotypes at low depth may lead to patterns of drift that are hard to distinguish from true biological signals.

To examine whether the GL method is robust to these issues, we introduced an additional three simulation scenarios in addition to the main simulation already described. See figure 1d for a summary of the differences between the four. In the main simulation (Uniform), discussed above, all populations have uniform depth 2 x, and there is no gene flow. In one alternative model (Migration), we add a pulse migration from population C into E 1000 generations before sampling time, with 50 % of the population replaced. In the two others, we keep the same tree topology, but change the simulated depths so that C and E are 15 x (C/E High), or so that all populations except C and E are 15 x (C/E Low). For clarity, in the below, we refer to the tree (i.e. the true topology for Uniform, C/E High, and C/E Low) as *G*_tree_, and the migration graph (used for Migration) as *G*_mig_.

For each of the additional simulation models, and for five independent replicates, we repeated the entire workflow from the main model to estimate drift trees with qpGraph and TreeMix. In each case, we fit the data on *G*_mig_ using qpGraph. We also fit the data using TreeMix with no allowed migration, which leads to inference of correct trees (or, for Migration, in some cases the tree with E and C as sister groups). The full trees for the first replicate of all models, normalised and unnormalised, are seen in figures S3 to S6.

In the tree model, we have a cline of increasing drift from populations A through G. Of course, this should be true regardless of the sequencing depth. However, the pulse in the Migration model will have the effect of breaking this cline at E. This is most easily seen in we focus on this part of the fitted tree, shown for the qpGraph fit in figure 1e (TreeMix in figure S7). Here we see that all four methods capture this break, although to various degrees of accuracy, and with various scaling of the drift. What is noteworthy, though, is that when calling genotypes, similar patterns emerge when the depth varies across populations (i.e. C/E High and C/E Low). In other words, differences in depth may cause a similar signal to gene flow.

To highlight that may in fact cause errors in inference, we proceeded to fit the *G*_mig_ model on all the data sets using qpGraph. (Overall drift inferred can be seen in figures S8 and S9). The inferred admixture proportion can be seen in figure 1f and shows that patterns of varying depth may in fact cause qpGraph to infer increased levels of migration. In particular, genotype calls with a frequency prior under the C/E models are inflated, with up to 15 % gene flow.

Moreover, we looked at whether qpGraph is able to fit the *f*_4_(*A, C*; *E, G*) statistic in the *G*_tree_ graph, corresponding to the amount of drift between where C splits off and where E splits. As expected, qpGraph always rejects the fit (i.e. reports a *z*-score greater than 2) of this *f*_4_ statistic in the Migration simulation when fitting *G*_tree_. Likewise, we would expect that this *f*_4_ statistic should be well-fitted in *G*_tree_ on the remaining three models, for which *G*_tree_ is in fact the true topology. This turns out to be true in all replicates for the true genotypes and the GL method, but it is false in multiple replicates when calling genotypes, whether using a frequency or uniform prior.

In summary, when calling genotypes with uneven depth, qpGraph frequency rejects that the topology is tree-like, even when the true topology is in fact a tree. In the absence of ground truth, this would lead to fitting a migration model, where migration may in fact be inferred.

### Hudson’s *F*_st_

As another example of a statistic we can get from the SFS with standard errors, we also calculate Hudson’s *F*_st_ [25] using sfs. This is shown for five replicates of the main simulation model in figure S10. Again, we find that *F*_st_ and 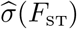 from the GL method closely recovers that of the true genotypes, while genotype calling biases the estimates downwards in an uneven manner across the tree.

### Relatedness

To illustrate the utility of derived statistics with standard errors on a real data set, we used whole genome sequencing data from 62 bush pigs and red river hogs, as well as a warthog sample used as outgroup. We refer to this data simply as the pig data set.

The samples in the pig data set stem from 13 distinct populations, which have names based on the country of sampling. The majority of the samples come from either Madagascar or Equatorial Guinea and have medium-low depth of coverage around 5 x. Many of the other populations are only represented by a single individual, typically at high depth. A summary of the data is shown in table S1. The full data set was sequenced and mapped as described in [26].

This data set is on the one hand intended to be typical of data that researchers working with low-depth data may encounter. At the same time, with most of the samples sequenced at medium-low depth, and broad variability in depth and sample size, it is also representative of a data set where we may worry about the effects of genotype calling.

Inference of relatedness may itself be of biological interest, but it is also often required in order to exclude samples from analyses assuming unrelated individuals. It is standard procedure to estimate kinship or relatedness from sequencing data, and a number of methods also exist for the case when genotype calling introduce bias [27, 28]. Moreover, statistics like R0, R1 [11], and King [29] can be calculated from the two-dimensional SFS between two individuals from sequencing to estimate relatedness [11], which provide a separate avenue for good estimates without genotype calling.

However, the variability of these estimates is generally not given, which may pose a challenge in practice. For example, say we are trying to exclude related individuals based on the King statistic. For context, the King kinship statistic is expected to be 0 for unrelated individuals from the same panmictic population, and positive for related pairs, with e.g. first-degree relatives expected on average to have a King kinship of 0.25. Suppose, then, that we are considering a pair with King 0.2, and our conclusion may well differ depending on whether the standard error is 0.02 or 0.2.

It is also important to know whether the standard error takes both the technical and the biological uncertainly into account. The latter consists of both linkage disequilibrium and the random recombination effects. If the block size used for the jackknife are large enough to extend beyond LD and IBD tracts then it takes the biological uncertain into account. Conversely, if the block size is small (e.g. a single base), then it only takes the technical uncertainly into account, which arise from the low depth and sequencing errors.

To estimate relatedness statistics for the pig dataset, we calculated genotype likelihoods for all 62 individuals using `angsd` and calculated the global and block spectra using `winsfs` with default settings for all intra-population pairs. Based on these, we calculated R0, R1, and King statistics with standard errors using 50 equally sized blocks. Of these, the King (figures 2a and S12) and R0 statistics (figure S11) are of general utility in diagnosing relatedness, whereas the R1 statistic (figure 2b) is mostly useful to distinguishing full siblings from parent-offspring pairs, where the other statistics lack power. We see a high level of background relatedness, especially in Madagascar, and some signs of population structure within some of the populations. We also see a number of pairs in the Madagascar and Uganda populations that appear to be first-degree relatives, likely split between parent-offspring pairs and full siblings. Moreover, the inclusion of standard errors allow us to judge which pairs are consistent with which relationships, given a particular threshold.

**Figure 2:**
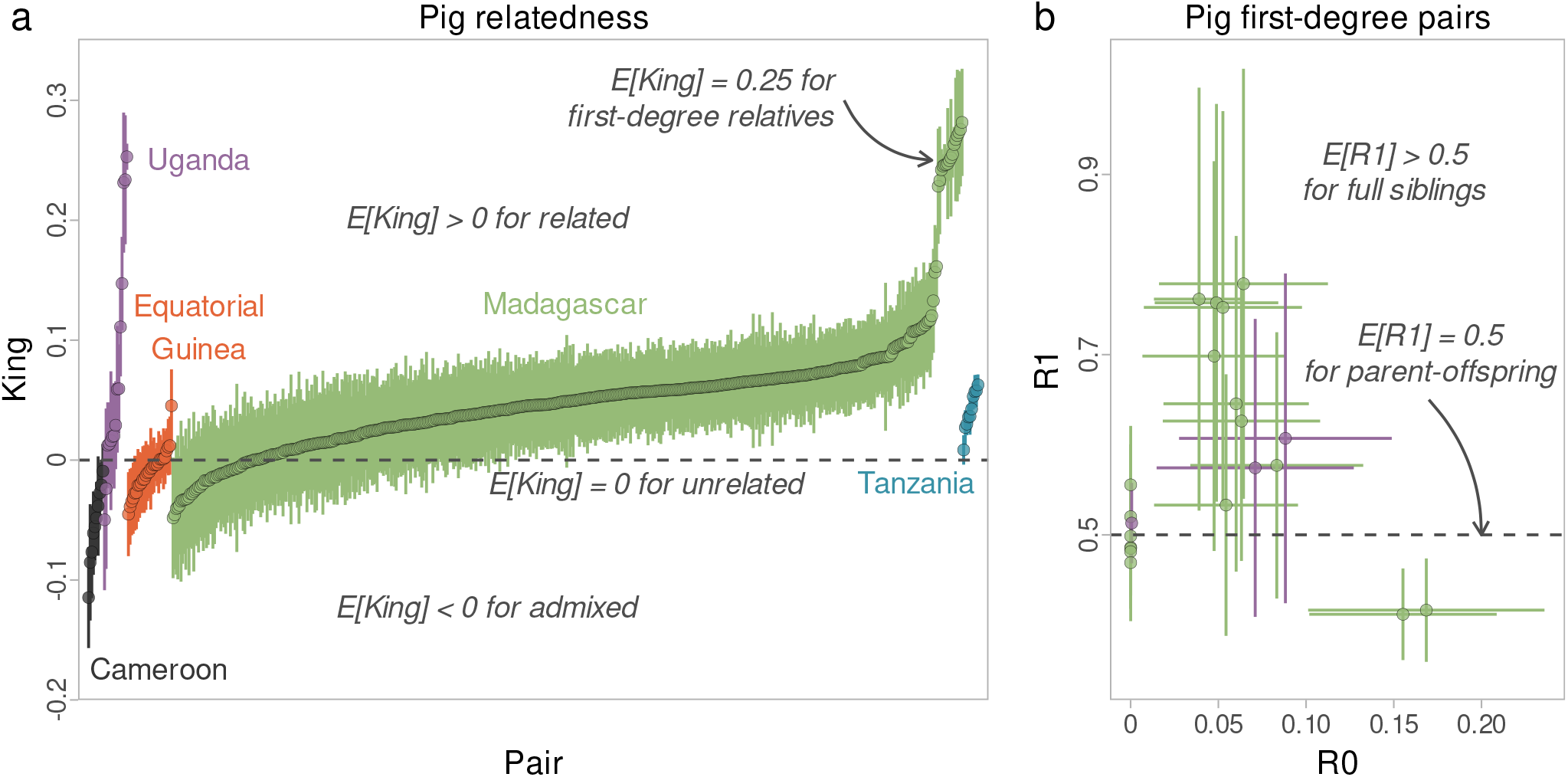
Relatedness statistics for the pig data set with italised annotations to aid interpretation (see [11] for details). **(a)**: King statistic *±*3 standard errors for all 559 intra-populations pairs of individuals from populations with at least three samples. **(b)**: R0 and R1 statistics *±*3 standard errors for all pairs with R1 greater than 0.4.

## Discussion

Overall, we have presented a framework for accurately computing a range of statistics with standard errors for low-depth whole-genome sequencing data. In particular, we have focused on how to accurately estimate *f*_2_ and the sample covariance *C* with standard errors, and how this enables accurate estimation of drift using the popular TreeMix and qpGraph tools.

In practice, we may often be more interested in inferring tree topologies with drift, as opposed to only fitting drift for a particular topology. Both TreeMix and the `find_graphs` function in ADMIXTOOLS2 offer this functionality and can be run using the methods described in this paper. However, finding the correct topology even with perfect genetic data is a harder problem than fitting drift for a particular topology, and is beyond the scope of this paper. We note that are many possible ways to explore the space of possible topologies for a set of populations, but where the patterns of drift are distorted, the resulting topology is likewise less likely to be correct. Therefore, we have confined ourselves to show that the bias in 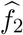 and *Ĉ* arising from genotype calling at low depth leads to such distortion instead of exploring all possible typologies. Fixing this distortion should always help downstream inference, regardless of what exact methods we employ.

We emphasise, though, that our approach to the estimation of standard errors targets the genome-scale setting. We compute this efficiently by estimating the SFS separately for each block of the genome. If we add those together we approximate the SFS for the genome, and based on our results this approximation give results very close to having perfect genotypes. However, it is unlikely to hold for SNP chips or exome data where we only have a limited number of sites. The flipside, of course, is that the need for such approximations is likewise lower with such data, where less care needs to be taken to stay within computational limits, and more straightforward methods can be applied.

At the intended genome scale, we have attempted to ensure that the current approach scales well. For example, computing 559 two-dimensional spectra and associated blocks for all intra-specific pig pairs was possible on a standard computing cluster overnight. This relies largely on previous work to optimise winsfs [9]. Finding the global SFS using winsfs is very efficient, and starting block estimates from the global estimate helps cut down time to convergence for the blocks. We might worry that doing so would make each block tend towards the global estimate, which would cause a downwards bias in the standard error estimates, but this does not appear to be an issue in practice. At the implementation level, care has also been taken to efficiently parallelise, to ensure cache locality, and to make use of vector registers where available.

As we have emphasised throughout, estimating standard errors on derived statistics for low-depth sequencing data have other uses than those emphasised here. An abundance of additional statistics can be derived from the SFS, including heterozygosity, the number of segregating sites and average pairwise differences (i.e., *π*) [30], average pairwise differences between populations (i.e., *d*_*XY*_) [31, 32], various estimators of *θ* and Tajima’s *D* [10]. Further, we can imagine running various SFS-based methods on the individual blocks for uncertainty estimates e.g. on demographic parameters with *∂a∂i* [33] or fastsimcoal [34, 35]. More generally, accurate and efficient block-wise estimates of the SFS or derived statistics may be useful, and might for instance form the basis for future work to detect introgression or selection by scanning statistics across the genome.

## Code and data availability

The blocked SFS estimation is available as part of the winsfs utility, available at github.com/malthesr/winsfs. The pig data set can be accessed via [26].

## Acknowledgements

AA, CW, and MSR are supported by the Independent Research Fund Denmark (grant numbers: 8021-00360B, 0135-00211B) and the University of Copenhagen through the Data+ initiative.

