## Supplementary material for "Inferring drift, genetic differentiation, and admixture graphs from low-depth sequencing data"

### Supplementary materials

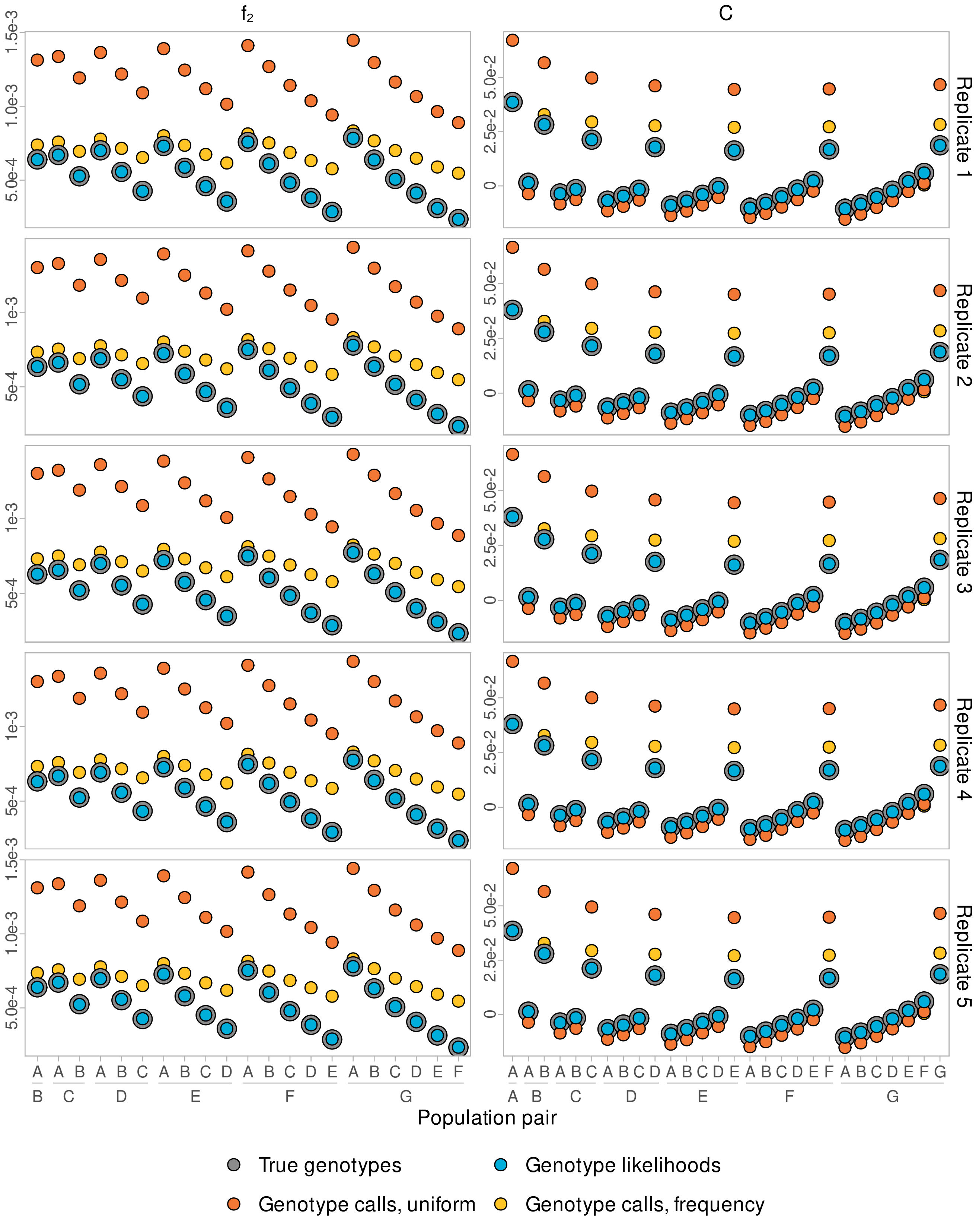


**Figure S1:** Estimates of $f_2$ and $C$ for five independently seeded replicates of the main simulation model.

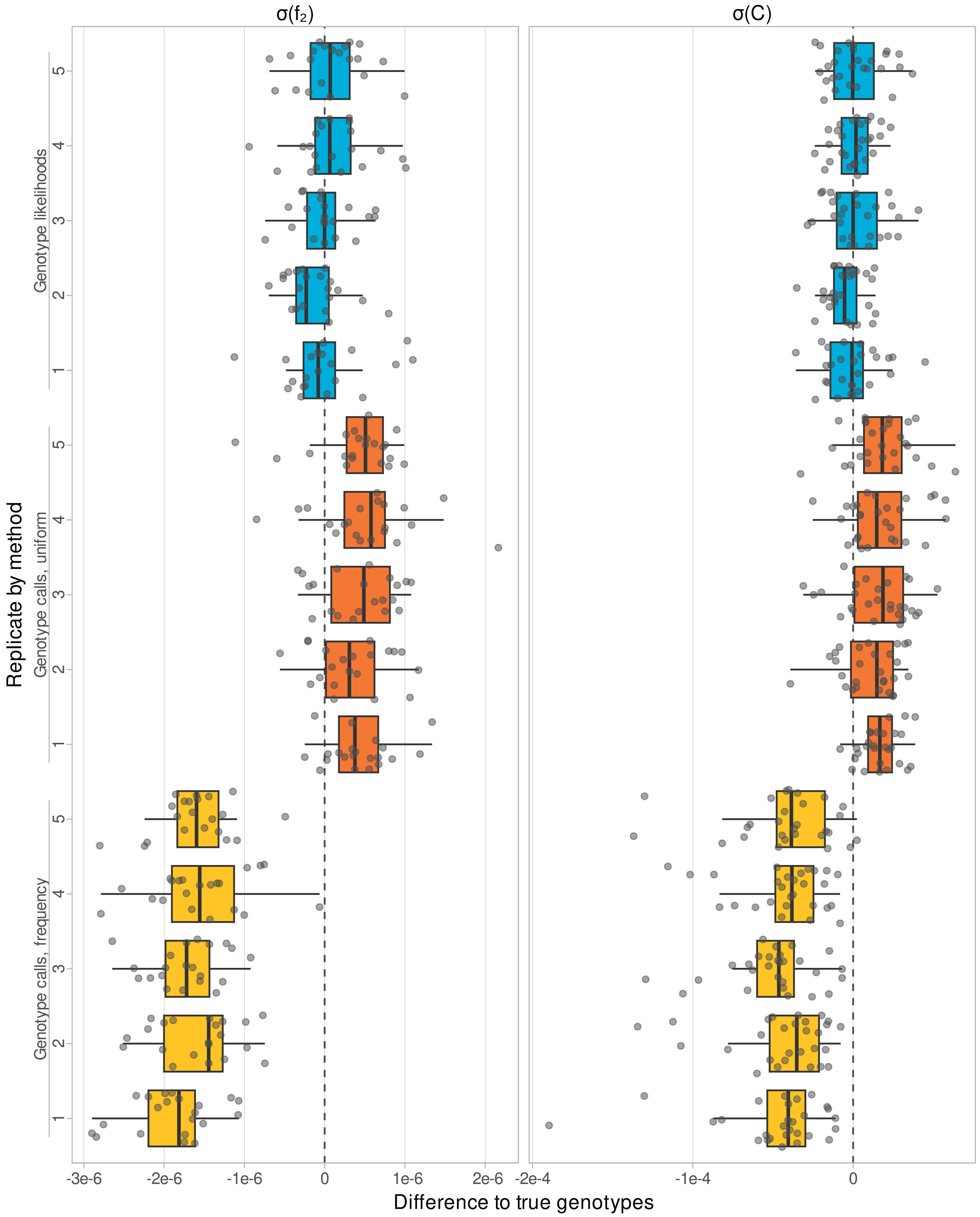


**Figure S2:** Estimates of $\sigma(f_2)$ and $\sigma(C)$ for five independently seeded replicates of the main simulation model. Compared to figure S1, the standard errors have slightly more noise, so these results are shown as difference to the true genotypes. The individual pairs are overlaid the boxplots as points with vertical jitter.

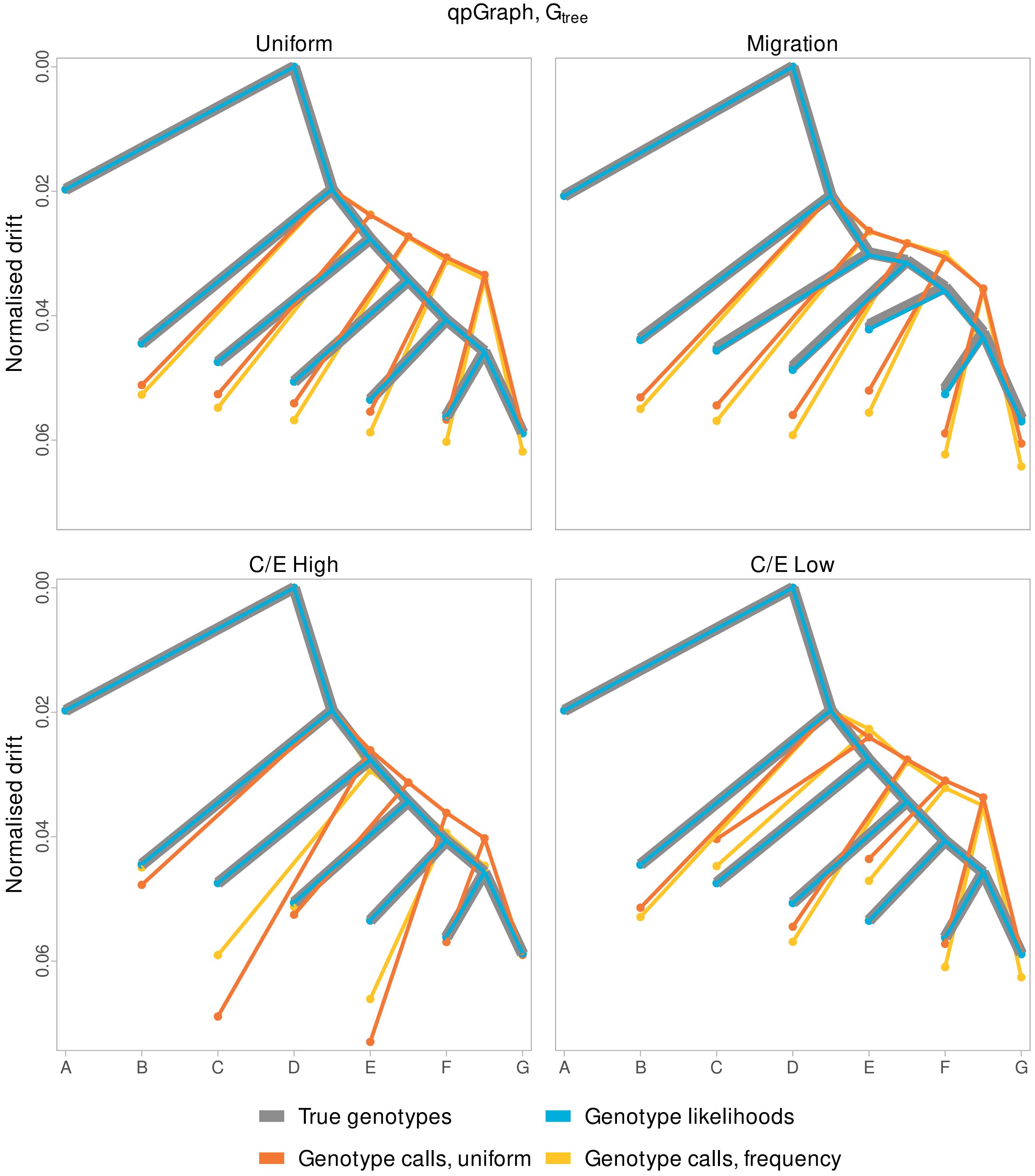


**Figure S3:** Normalised drift inferred by qpGraph for all four simulation models on the tree model $G_{\text{tree}}$. The inferred drift is normalised by the branch length to the outgroup A.

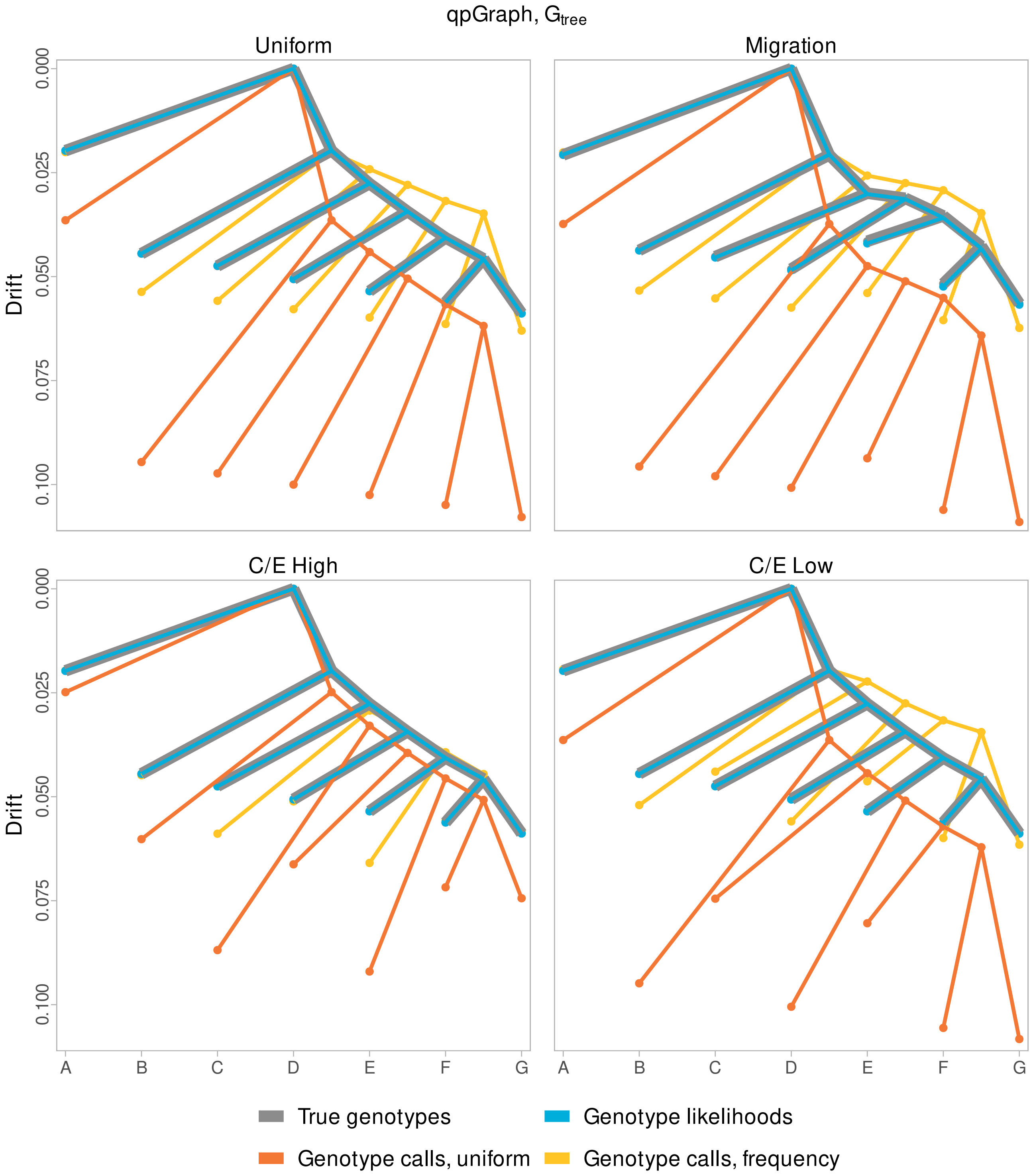


**Figure S4:** Unnormalised drift inferred by qpGraph for all four simulation models on the tree model $G_{\text{tree}}$.

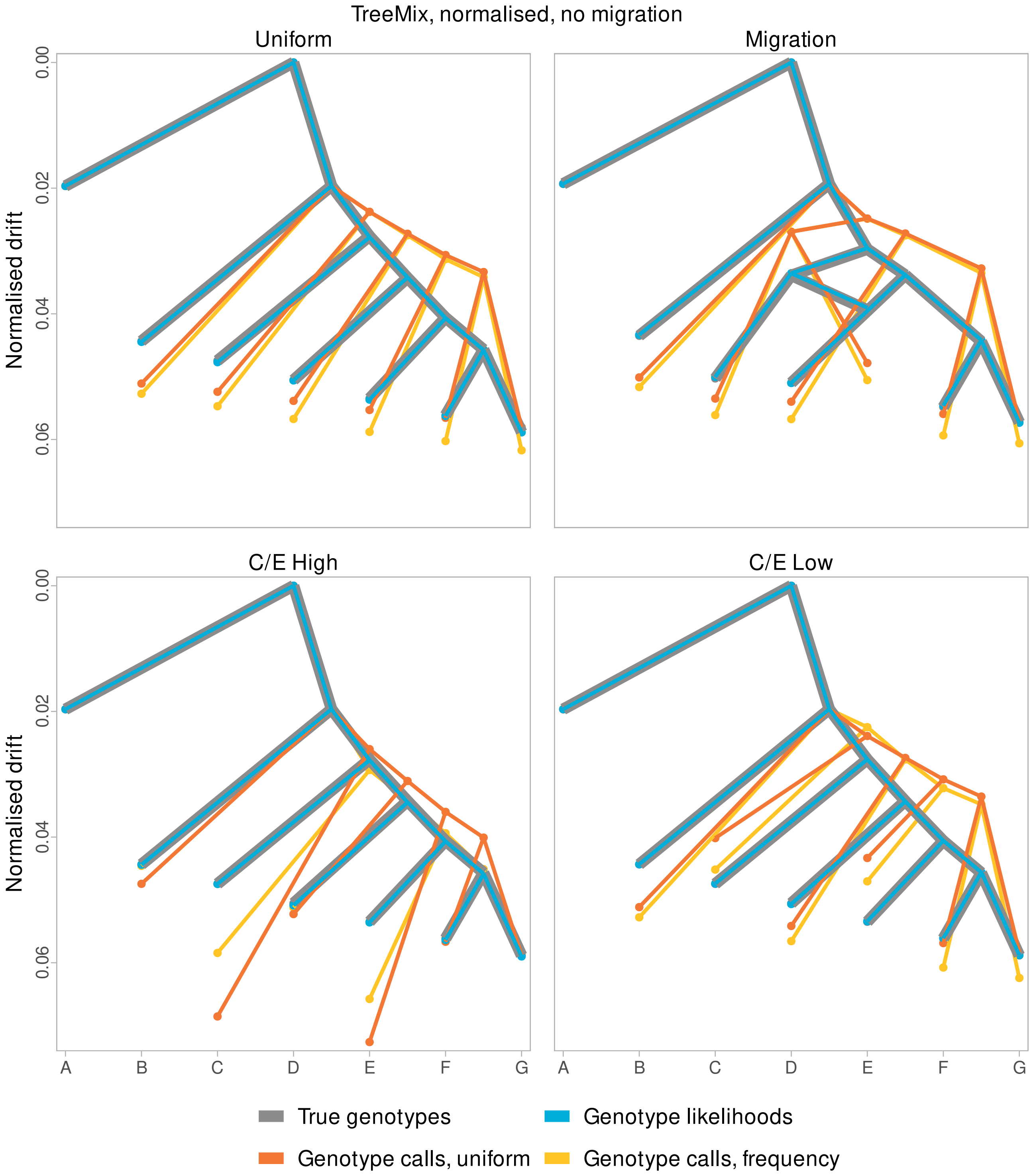


**Figure S5:** Normalised drift inferred by TreeMix for all four simulation models with no migration allowed. The inferred drift is normalised by the branch length to the outgroup A.

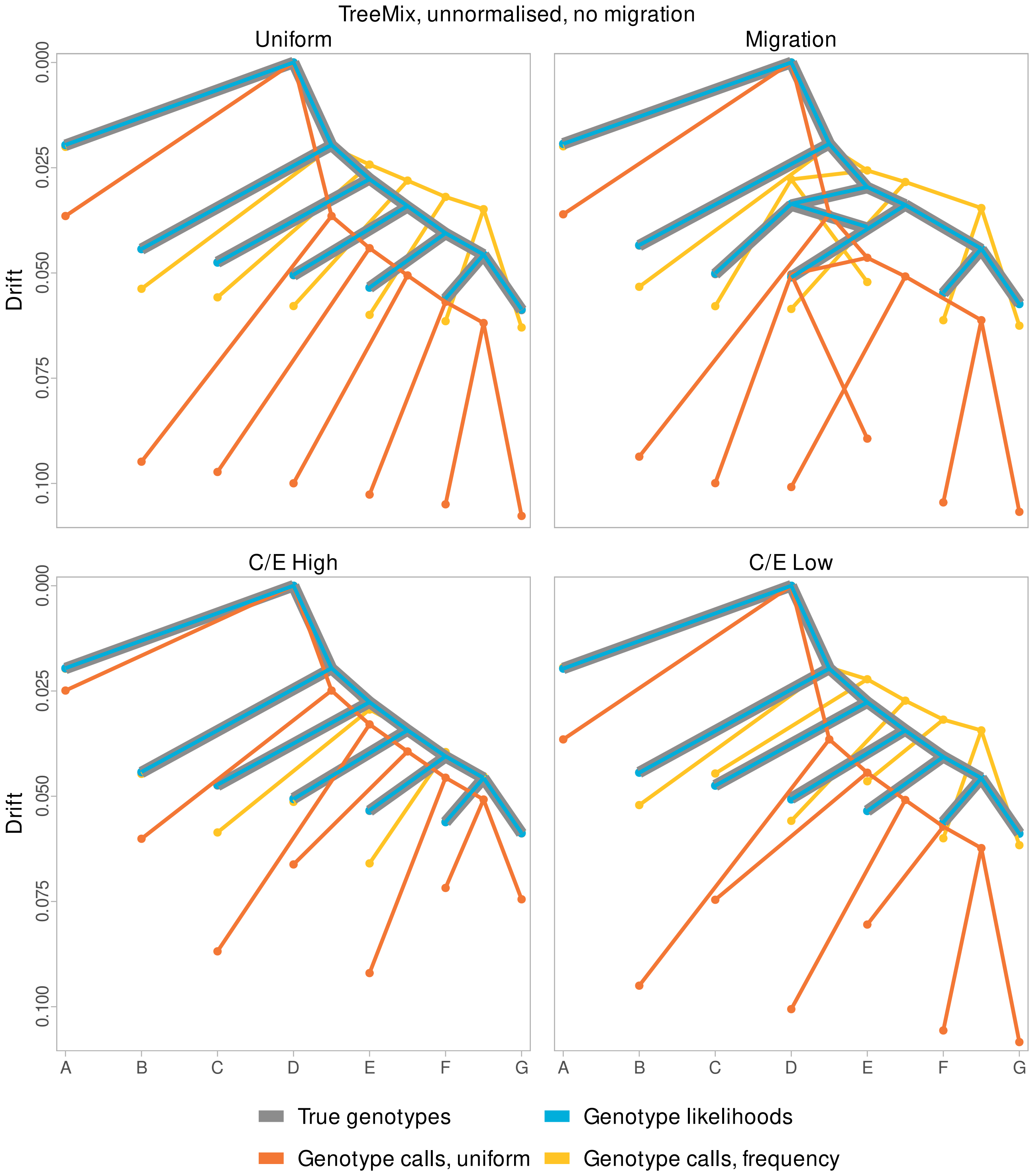


**Figure S6:** Unnormalised drift inferred by TreeMix for all four simulation models with no migration allowed.

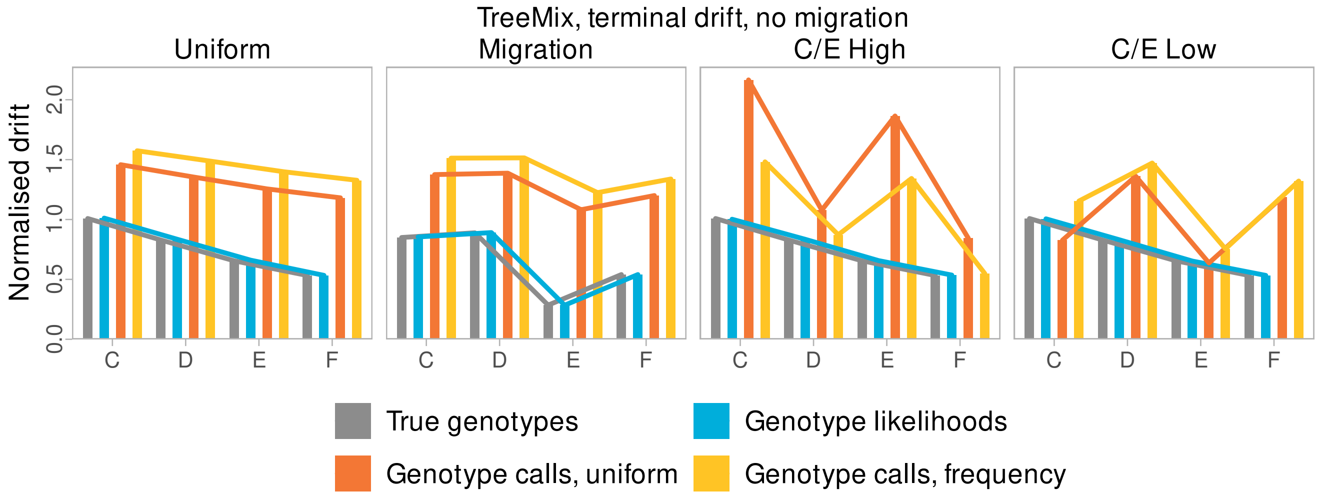


**Figure S7:** Normalised terminal drift estimated by Treemix with no migration allowed for populations C, D, E, F including alternative models. Drift between internal nodes not to scale.

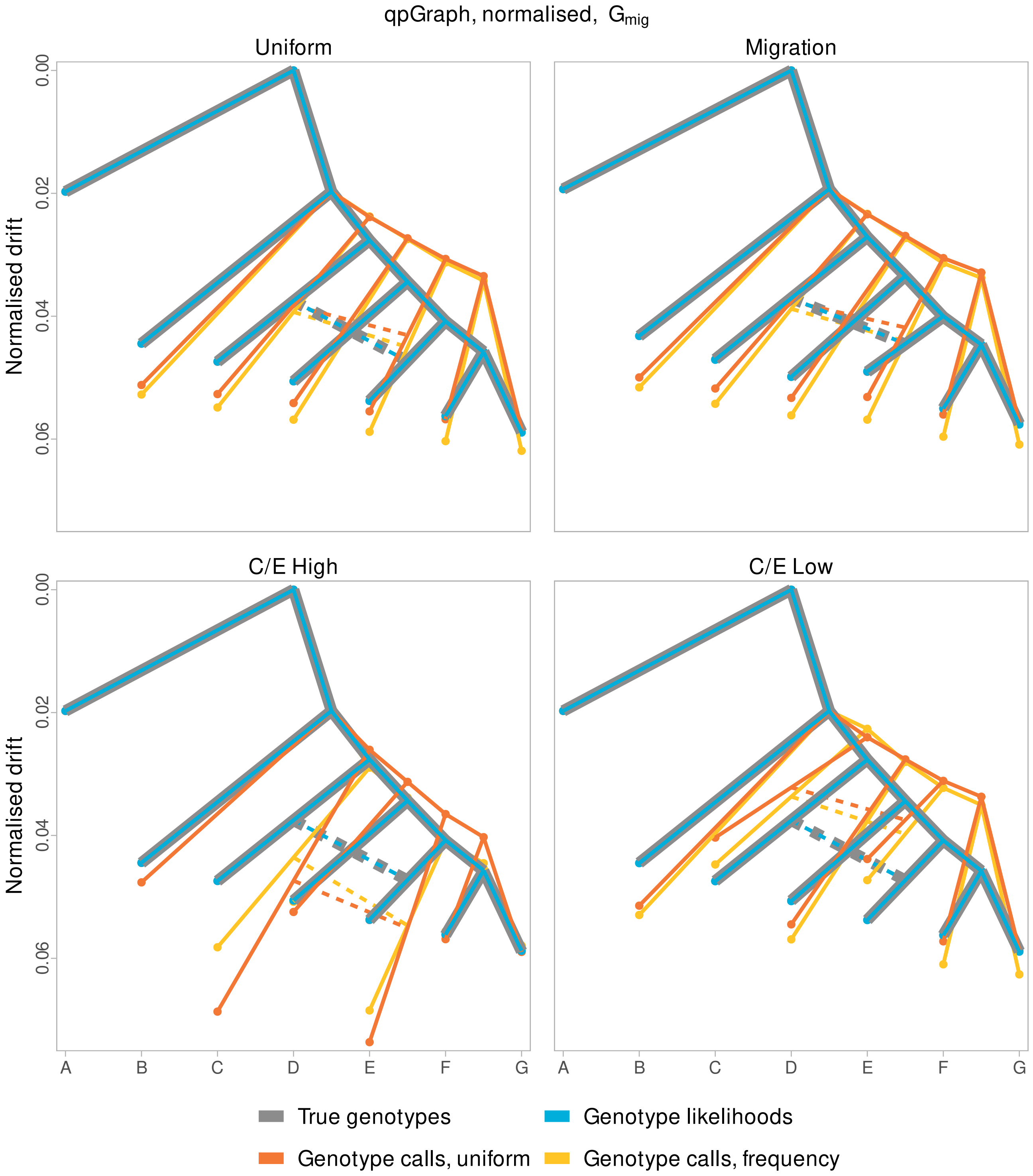


**Figure S8:** Normalised drift inferred by qpGraph for all four simulation models on the tree model $G_{\mathrm{mig}}$. The inferred drift is normalised by the branch length to the outgroup A. Placement of migration on edges to C and E is arbitrary.

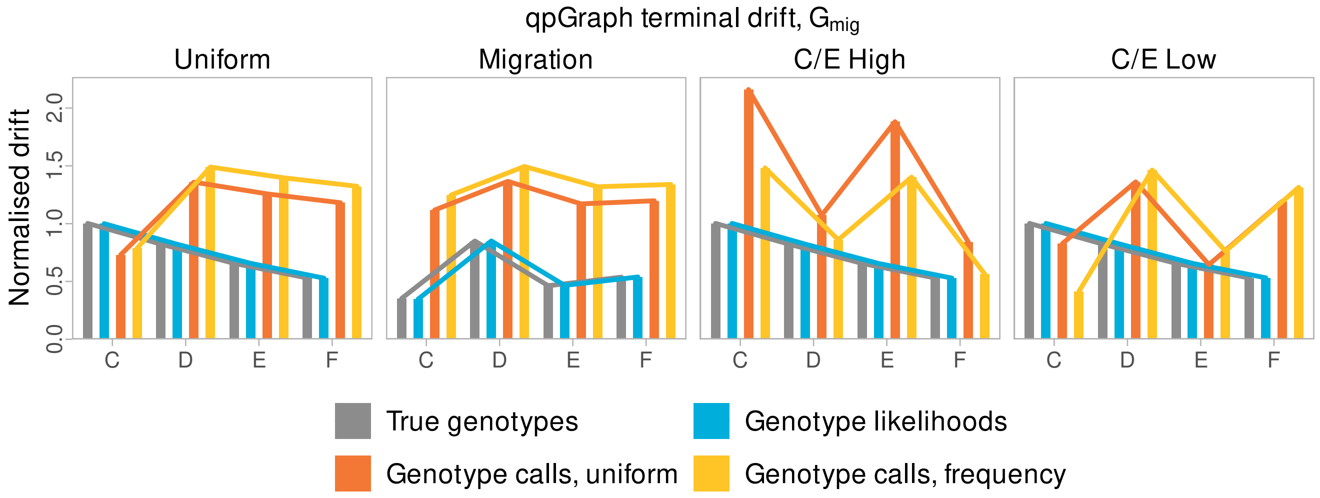


**Figure S9:** Normalised terminal drift estimated by qpGraph for populations C, D, E, F including alternative models, fitted on graph that includes migration from C to E. Drift between internal nodes not to scale.

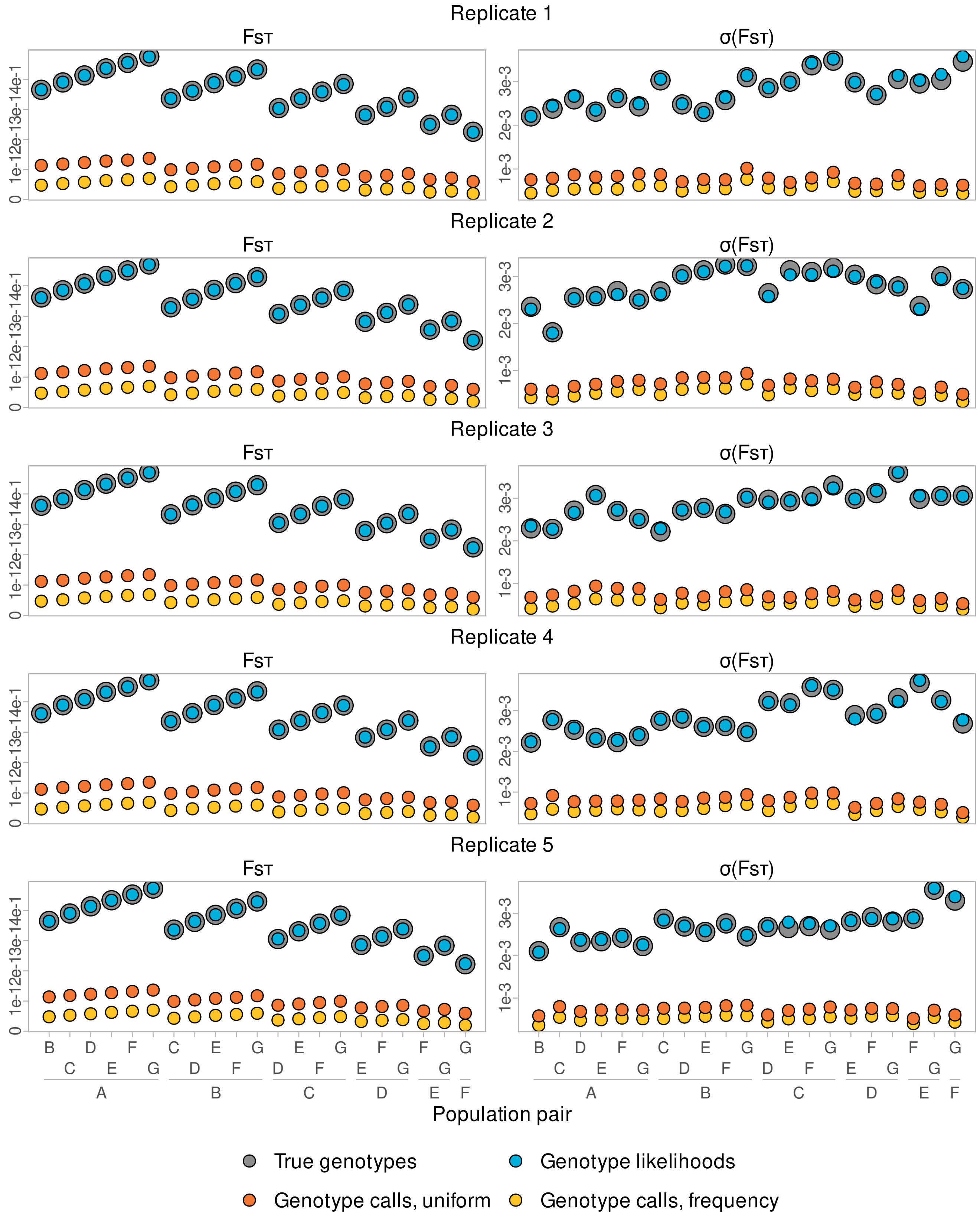


**Figure S10:** Estimates of Hudson's $F_{ST}$ with standard errors for five replicates of the main simulation model.

| Population name | Sample size | Median depth | Sample size after filtering | Median depth after filtering |
|---|---|---|---|---|
| Madagascar | 32 | 5.2 x | 21 | 5.2 x |
| Equatorial Guinea | 8 | 4.0 x | 8 | 4.0 x |
| Uganda | 6 | 34.0 x | 4 | 34.0 x |
| Tanzania | 5 | 5.0 x | 5 | 5.0 x |
| Cameroon | 5 | 4.0 x | 5 | 4.0 x |
| DRC | 2 | 3.8 x | 2 | 3.8 x |
| Ethiopia | 2 | 71.6 x | 2 | 71.6 x |
| Nigeria | 2 | 64.8 x | 2 | 64.8 x |
| Ghana | 1 | 17.9 x | 1 | 17.9 x |
| Gabon | 1 | 13.9 x | 1 | 13.9 x |
| Togo | 1 | 15.1 x | 1 | 15.1 x |
| South Africa | 1 | 17.0 x | 1 | 17.0 x |
| Zimbabwe | 1 | 19.7 x | 1 | 19.7 x |
| Warthog | 1 | 60.9 x | 1 | 60.9 x |

**Table S1:** Overview of the pig data set by populations. The sample size and median depth is shown both before and after filtering out related individuals.

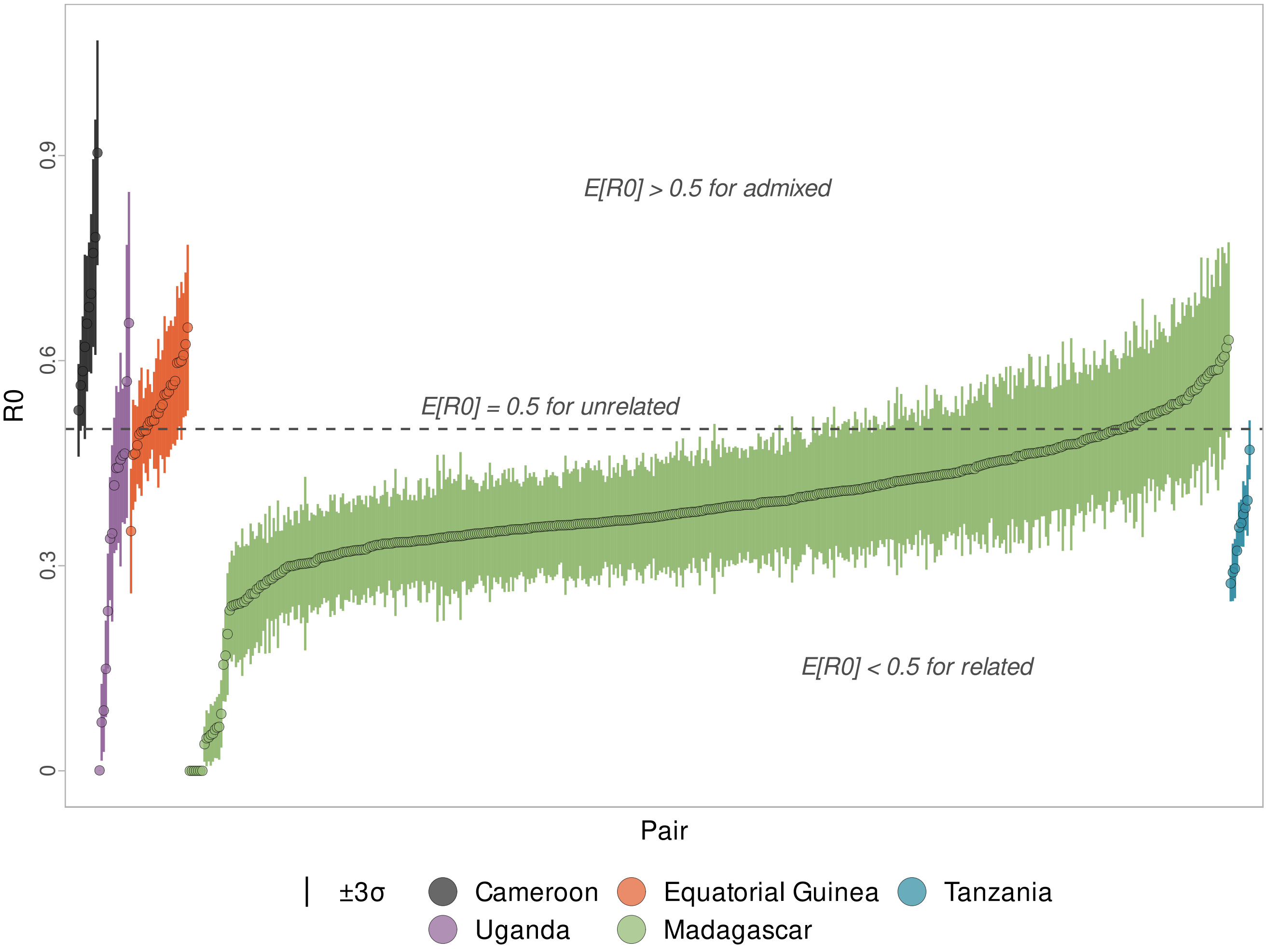


**Figure S11:** R0 statistic ±3 standard errors for all 559 intra-populations pairs of individuals from populations with at least three samples. Annotations are shown to give some context for the expectation of the statistic under various scenarios. Note that $E[R_0] < 0.5$ for related individuals under the assumption that they are not admixed and that the pair is from the same population.

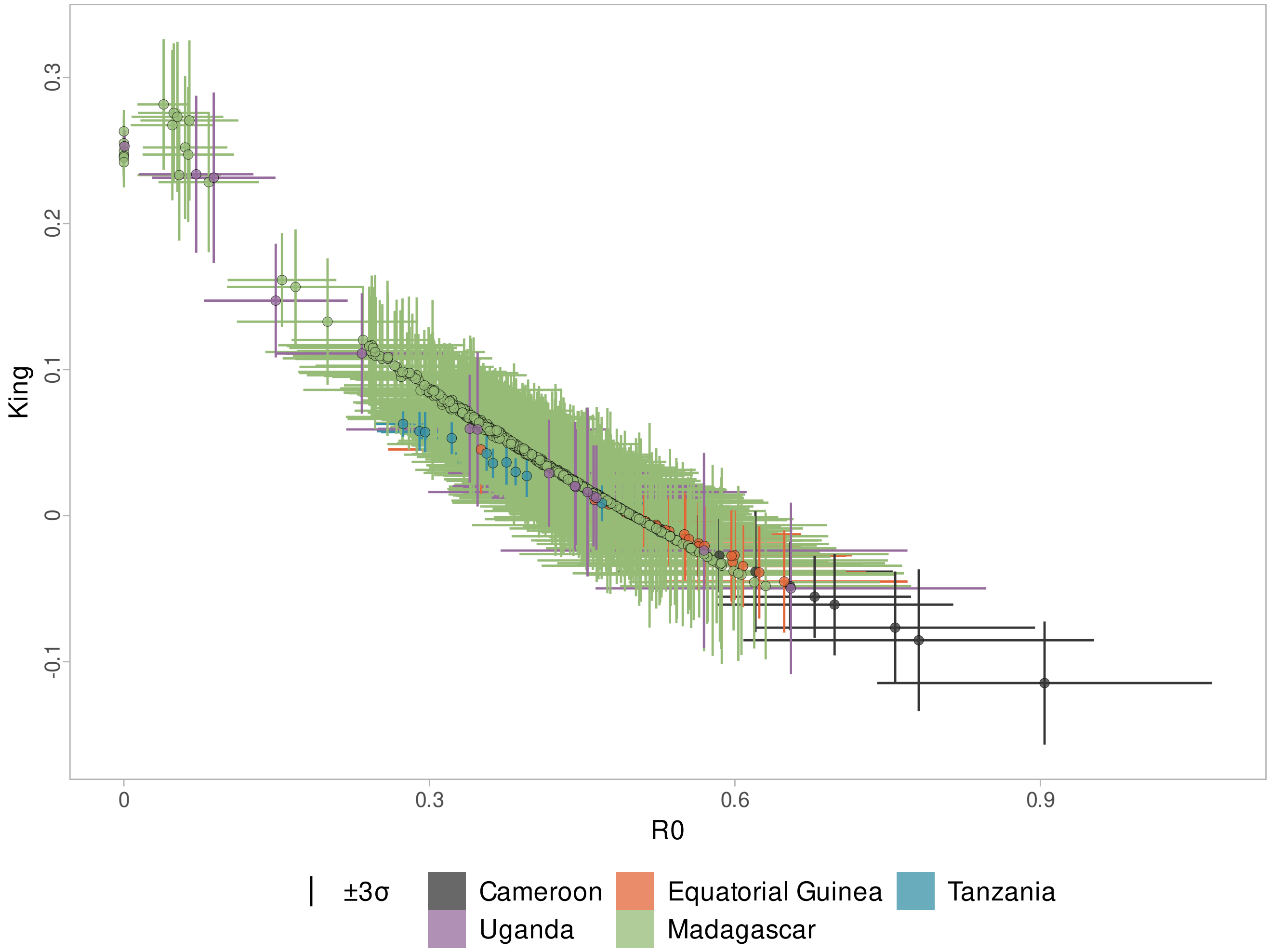


**Figure S12:** R0/King statistics ±3 standard errors for all 559 intra-populations pairs of individuals from populations with at least three samples. Note that the parent offspring will have $E[R_0] = 0$ while the full siblings will have a $E[R_0] > 0$.
